# Directionality bias underpins divergent spatiotemporal progression of Alzheimer-related tauopathy in mouse models

**DOI:** 10.1101/2024.06.04.597478

**Authors:** Justin Torok, Christopher Mezias, Ashish Raj

## Abstract

Mounting evidence implicates trans-synaptic connectome-based spread as a shared mechanism behind different tauopathic conditions, yet also suggests there is divergent spatiotemporal progression between them. A potential parsimonious explanation for this apparent contradiction could be that different conditions incur differential rates and directional biases in tau transmission along fiber tracts. In this meta-analysis we closely examined this hypothesis and quantitatively tested it using spatiotemporal tau pathology patterns from 11 distinct models across 4 experimental studies. For this purpose, we extended a network-based spread model by incorporating net directionality along the connectome. Our data unambiguously supports the directional transmission hypothesis. First, retrograde bias is an unambiguously better predictor of tau progression than anterograde bias. Second, while spread exhibits retrograde character, our best-fitting biophysical models incorporate the mixed effects of both retrograde- and anterograde-directed spread, with notable tau-strain-specific differences. We also found a nontrivial association between directionality bias and tau strain aggressiveness, with more virulent strains exhibiting less retrograde character. Taken together, our study implicates directional transmission bias in tau transmission along fiber tracts as a general feature of tauopathy spread and a strong candidate explanation for the diversity of spatiotemporal tau progression between conditions. This simple and parsimonious mechanism may potentially fill a critical gap in our knowledge of the spatiotemporal ramification of divergent tauopathies.

## Introduction

It is well known that the spatiotemporal pattern of Alzheimer’s disease (AD) pathology development is characterized by stereotyped progression. Braak & Braak were the first to describe the staging of hallmark neurofibrillary tangles (NFT) of misfolded tau protein: these are first detected in the entorhinal cortex (EC) and locus coeruleus (LC), followed by limbic areas and then neocortical areas while sparing primary sensorimotor cortices ***Braak and Braak (1991)***. Identifying the causal mechanisms that govern why tau follows this characteristic spreading pattern in AD is a longstanding goal of neurodegenerative disease research. Progress on this front has rapidly advanced with the discovery that hyperphosphorylated tau undergoes intercellular spread following axonal transport and trans-synaptic transmission ***Ahmed et al. (2014)***; ***Clavaguera et al. (2009)***; ***Holmes et al. (2014)***; ***Tai et al. (2014)***).

However, key aspects of the transmission process remain unknown, especially whether axonal tau travels bidirectionally or preferentially in either in the same direction (anterograde) or in the reverse direction (retrograde) with respect to the neuron’s polarity. Similarly, it is not known whether the trans-synaptic transmission occurs from post-to pre-synapse, post-to pre-synapse, or some mixture of the two. Much phenomenological evidence and descriptive studies are now available to support both anterograde and retrograde transmission ***Wu et al. (2013)***; ***Langer Horvat et al. (2023)***. However, these studies do not provide a statistically rigorous support to net directionality across multiple disease models, nor do they allow inference of the net directional bias in those cases where it may not be either fully anterograde or retrograde. Most importantly, current methods are unable to tie directional bias to more fundamental processes in the aggregation and transmission of tau along axons, which may ultimately be required to explain how different strains of tau exhibit such strikingly different spatial patterns over time.

The purpose of this study is to assess whole-brain and unbiased statistical support for net directionality gleaned from integrated large-scale meta-analysis of all available tauopathic data measured in our chosen model system - transgenic mouse models of AD tauopathy. We hope to move beyond current evidence that has understandably come from descriptive, hypothesis-driven mechanistic experimental bench or animal studies. These studies, based on selected brain structures or selected tau conformers or strains, are difficult to generalize and frequently produce conflicting results. Instead we aim for “big picture” and generalizable statistical evidence that together supports the emergence of directional bias in tau transmission. We achieve these goals by analysing histological tau data from 4 individual studies comprising 11 experimental conditions, all of which use P301S-tau-expressing mice***Boluda et al. (2015)***; ***Hurtado et al. (2010)***; ***Iba et al. (2013)***; ***Kaufman et al. (2016)***, to perform a meta-analysis of mouse tauopathy at a whole-brain level in the context of network spread of tau.

We approached this task in two distinct ways. First, we employed statistical and graph theoretical techniques to generate evidence for directional bias in tau spread, and report that indeed empirical data cannot be explained without introducing directionality of transmission. Second, we used novel mathematical models of tau network spread, which enabled us to explore directional bias in the context of other key pathological processes, such as accumulation of tau. In particular, we investigated whether intrinsic net directional bias in trans-synaptic spread can better recapitulate empirical pathology data. For this purpose, we used as our base model our previous mathematical network spread models of pathology spread along anatomical connectivity successfully recapitulate the spatiotemporal regional volumetric loss in patients’ brains. These models, developed by our group ***Anand et al. (2022)***; ***Raj et al. (2012, 2015)***; ***Mezias et al. (2017)*** and others ***Iturria-Medina (2013)***; ***Weickenmeier et al. (2018)***; ***Bertsch et al. (2023)***, show great promise in predicting later disease patterns from early timepoints.

The specific mechanistic model we employed is an extension of the NexIS model of pathology spread ***Anand et al. (2022)***, expanded to incorporate directionally biased transmission in both anterograde (pre-to post-synaptic) or retrograde (post-to pre-synaptic) directions. This model has previously been shown to more accurately recapitulate the longitudinal progression of tauopathy in mouse models than prior network transmission models ***Raj et al. (2015)***. We call this model *NexIS:dir*. The model was simulated on the mesoscale connectome of the mouse, utilizing the Allen Mouse Brain Connectivity Atlas (AMBCA), which not only provides connection strengths between region-pairs but also the directionality of those connections ***Oh et al. (2014)***. This model-based method of studying connectivity and directionality has the advantage of not depending on transmission along a single projection, but incorporates directional transmission across the entire brain network.

As we show below, this approach allowed us to clearly demonstrate that retrograde spread bias is a characteristic and necessary feature of tau progression in many experimental models. We also showed that net directional bias has a dramatic effect on the spatial patterns of tau deposition over time. Our study therefore has the potential to explain strain-specific differences in spatial patterns of tau deposition.

## Results

### Analysis of directed connectivity graph indicates retrograde preference of tau migration

Previous network modeling has demonstrated that connectivity with afflicted regions drives subsequent spatiotemporal tau progression, but was limited to undirected connectomes ***Mezias et al. (2017)***; ***Raj et al. (2012, 2015)***; ***Zhou et al. (2012)***; ***Weickenmeier et al. (2018)***; ***Iturria-Medina (2013)***. This mimics a diffusion process whereby the direction of tau spread between region pairs is dictated only by the concentration difference between them. However, migration of tau along axonal projections is known to have both diffusive and active transport components ***Scholz and Mandelkow (2014)***, where molecular motors traffic cellular cargoes, including tau, asymmetrically between the soma and the axon terminal (**Figure 1A**). At a network level, this manifests as a *directionality bias*, with tau preferentially migrating in the *anterograde* (from presynaptic to postsynaptic regions, following the polarity of the projections) or *retrograde* (against the polarity of the projections) (**Figure 1B**). Exploring directionality requires the use of a *directed* connectome, which has been previously determined using viral tracing methods in wild-type mice by the Allen Institute for Brain Science (AIBS) ***Oh et al. (2014)***; here, we use a 426-region, bilateral version of the Allen Mouse Brain Connectome Atlas (AMBCA) (**Figure 1C**). This makes mouse models of tauopathy an ideal substrate for exploring the question of directional bias in tau spread. For this study, we utilized 11 distinct mouse models, whose descriptions can be found in **Table S1**). Of particular note is that 10 of 11 studies had the same genetic background (PS19), and therefore any differences between studies’ tau pathology can be attributed to differences in seeding location and the molecular properties of the tau strain injected.

**Figure 1.**
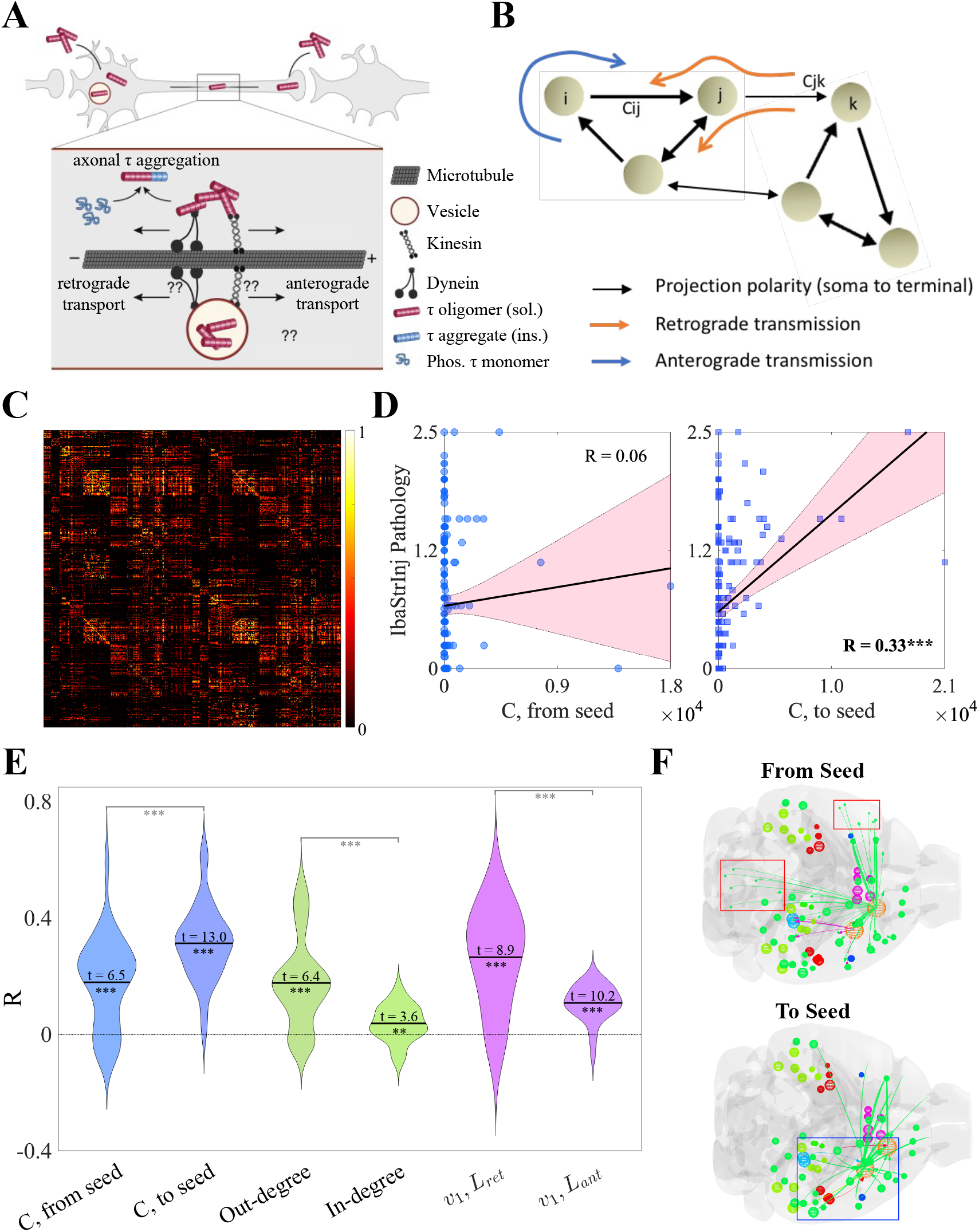
Tau exhibits a network-level directional preference. **A. & B**. Schematics showing the etiology of directional spread bias at the microscopic and network levels. Inside of axons, pathological tau can migrate by passive diffusion or through energy-dependent directed transport either in the *anterograde* (parallel to axon polarity) or *retrograde* (antiparallel to axon polarity) directions (**A**). At a network level, this manifests as an *directionally biased* flow along the directed connectome **B**. By convention, *c*_*ij*_ indicates a connection originating in region *i* and terminating in region *j*, therefore a net flow from *i* to *j* along *c*_*ij*_ would be considered to be *anterograde-biased*. **C**. The Allen Mouse Brain Connectome Atlas (AMBCA) ***Oh et al. (2014)*** visualized as a heatmap. **D**. Scatterplots showing the associations between the regional end-timepoint (9 MPI) pathology in the IbaStrInj ***Iba et al. (2013)*** experiment (see **Table S1**) and the average connectivity from (*left*) and (*right*) seeded regions CP and MOp. Tau shows a highly significant association with incoming but not outgoing connectivity. **E**. Violin plots showing the associations between tau pathology across all studies and time points and three pairs of region-level graph metrics; from right to left: outgoing and incoming connectivity to seed, out- and in-degree, and the first eigenvectors (*ν*_1_) of *L*_*ret*_ and *L*_*ant*_ (see **Materials and Methods**). One-sample t-statistics were calculated for each metric and two-sample t-statistics were calculated for each metric pair. All t-statistics were highly significant. **F**. Glass brain visualizations of end-timepoint IbaStrInj pathology and connectivity from (*top*) and to (*bottom*) seed regions (orange spheres). The low association between outgoing seed connectivity is in part driven by strong contralateral telencephalic and ipsilateral hindbrain projections (red boxes), which do not exhibit significant tau pathology. By contrast, the seeded regions predominantly receive connectivity from ipsilateral forebrain regions, which do exhibit pronounced tau pathology (blue box). MPI – months post injection; CP – caudoputamen; MOp – primary motor cortex. * – *p* < 0.05; ** – *p* < 0.01; *** – *p* < 0.001.

We first explored directional bias in a model-free way. **Figure 1D** shows the associations between end-timepoint tau pathology for the “IbaStrInj” mouse model ***Iba et al. (2013)***, which was quantified at 9 months post injection (MPI), and the connectivity from (*left panel*) and to (*right panel*) the seeded regions (in this case, the caudoputamen and primary motor cortex). We observed a strongly statistically significant association with respect to incoming connectivity to the seeded regions (Pearson’s *R* = 0.33, *p* < 0.001), but not outgoing connectivity (*R* = 0.06, n.s.). Because preferential spread along incoming connections indicates that tau is migrating from postsynaptic to presynaptic regions, these results suggest that the IbaStrInj mouse model exhibits a net retrograde directionality bias.

We found that this appears to be a general feature of mouse tauopathy for the 11 models we explored, albeit with significant heterogeneity between studies and timepoints, as assessed by three different sets of graph metrics: connectivity from/to seed, out- and in-degree, and the first eigenvectors of the retrograde and anterograde Laplacian matrices (*L*_*ret*_ and *L*_*ant*_, respectively; see also **Materials and Methods**) (**Figure 1E**). In graph theory the first eigenvector of the directional Laplacian is considered a “network sink” of directionally biased spread. While the distributions of *R* values were all significantly above 0 by one-sample t-test following Fisher’s *R*-to-*z* transformation (all *p* < 0.01), we found statistically significant differences within each pair of metrics. Specifically, associations with tau pathology were greater for incoming rather than outgoing connectivity with seeded regions; for out-degree rather than in-degree; and for *ν*_1_ of *L*_*ret*_ rather than *ν*_1_ of *L*_*ant*_ (all *p* < 0.001 by two-sample t-test). Examining differences in associations with *ν*_1_ is particularly instruc-tive, as this is the eigenmode of *L* that should be most associated with tau spread over slow time scales. *L*_*ret*_ and *L*_*ant*_ were constructed such that an outflux of tau from region *i* to region *j* occurs along connections *c*_*ji*_ (retrograde) and *c*_*ij*_ (anterograde), respectively (see **Materials and Methods**). Therefore, the fact that *ν*_1_ of *L*_*ret*_ has a significantly stronger association with tau pathology suggests that retrograde-biased spread dominates.

To examine these effects qualitatively, we returned to the end-timepoint pathology of the IbaStrInj mouse model and utilized a glass brain visualization of regional tau densities alongside the top 10% of connections from and to seeded regions (**Figure 1F**). At early stages, there is a net outflux of tau from seeded regions into connected regions. If that outflux tends to occur along outgoing connections (i.e., *c*_*ij*_, where *i* is a seeded region), then spread is anterograde-biased; the reverse is true for retrograde. For IbaStrInj, we observed that many of the strongest projections from the primary motor cortex (MOp, orange sphere) terminate in the contralateral telencephelon and ipsilateral hindbrain (*top panel*, red boxes), where this mouse model exhibits low pathology, as well as ipsilateral telencephalic regions, where pathology is higher. By contrast, incoming connectivity to both the MOp and caudoputamen (CP) is largely ipsilateral and originates in forebrain regions that exhibit high tau density (**Figure 1F**, *bottom panel*, blue box). Because these results indicate a preferential outflux from seed along projections *c*_*ji*_ rather than *c*_*ij*_, these results suggest that there is a retrograde rather than anterograde preference in mouse tauopathy.

### NexIS:dir modeling of tau pathology in IbaStrInj demonstrates moderately but not wholly retrograde spread

To model the spread of tau more directly, we extended the previously explored NexIS model ***Anand et al. (2022)*** into NexIS:dir, designed for directed connectomes, and introduces a new directionality bias parameter *s* that weights the spread along *L*_*ret*_ versus *L*_*ant*_, with *s* = 1 indicating purely retrograde spread, *s* = 0 indicating purely anterograde spread, and *s* = 0.5 indicating nondirectional spread, which is equivalent to spread along the symmetrized connectome 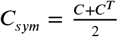 (see **Materials and Methods**). The other parameters in the model are: *α*, the net aggregation or clearance rate of tau, and *β*, its rate of diffusion into the connectome, i.e. spread rate. We postulated that, while spread may be net retrograde (**Figure 1D–F**), the best model of tau progression would require capturing intermediate directionality biases through *s*.

**Figure 2** shows glass brain representations of IbaStrInj pathology at 1, 3, and 9 MPI as well as the predictions of four different NexIS:dir models: 1) NexIS:fit-s, where *s* was fit alongside the accumulation (*α*) and spread (*β*) parameters; 2) NexIS:ret, where *s* was fixed to 1; 3) NexIS:ant, where *s* was fixed to 0; and 4) NexIS:nd, where *s* was fixed to 0.5. All model parameters were optimized utilizing all timepoints together (see **Materials and Methods**). We observed that the rapid dissemination of tau between 0 and 1 MPI in this mouse model was poorly captured by all four models. However, by the last quantified timepoint, NexIS:fit-s captured the highest number of relevant features in tau spread relative to all other models. According to this model, spread was predominantly, but not completely, retrograde, with an optimal *s* value of 0.78. Each of the other models where *s* was fixed underperformed in different ways. NexIS:ant, the worst model, could not fit the spread process at all and predicted that tau pathology remained in the seeded regions. NexIS:ret did capture the spread of tau to lateral neocortical regions observed in this mouse model, but showed little spread elsewhere. NexIS:nd was able to accurately predict the spread of tau to neocortical and forebrain subcortical regions; however, it overpredicted pathology in the ipsilateral and contralateral CP as well as the midbrain. NexIS:fit-s demonstrated a net decrease in pathology in the CP as well as spreading throughout lateral forebrain regions, with an excellent overall fit across all timepoints (*R* = 0.56, *p* < 0.001). Therefore, we conclude that, in order to truly predict the spatiotemporal deposition of tau in IbaStrInj, we require the precise quantification of the proportion of directionality bias through *s*.

**Figure 2.**
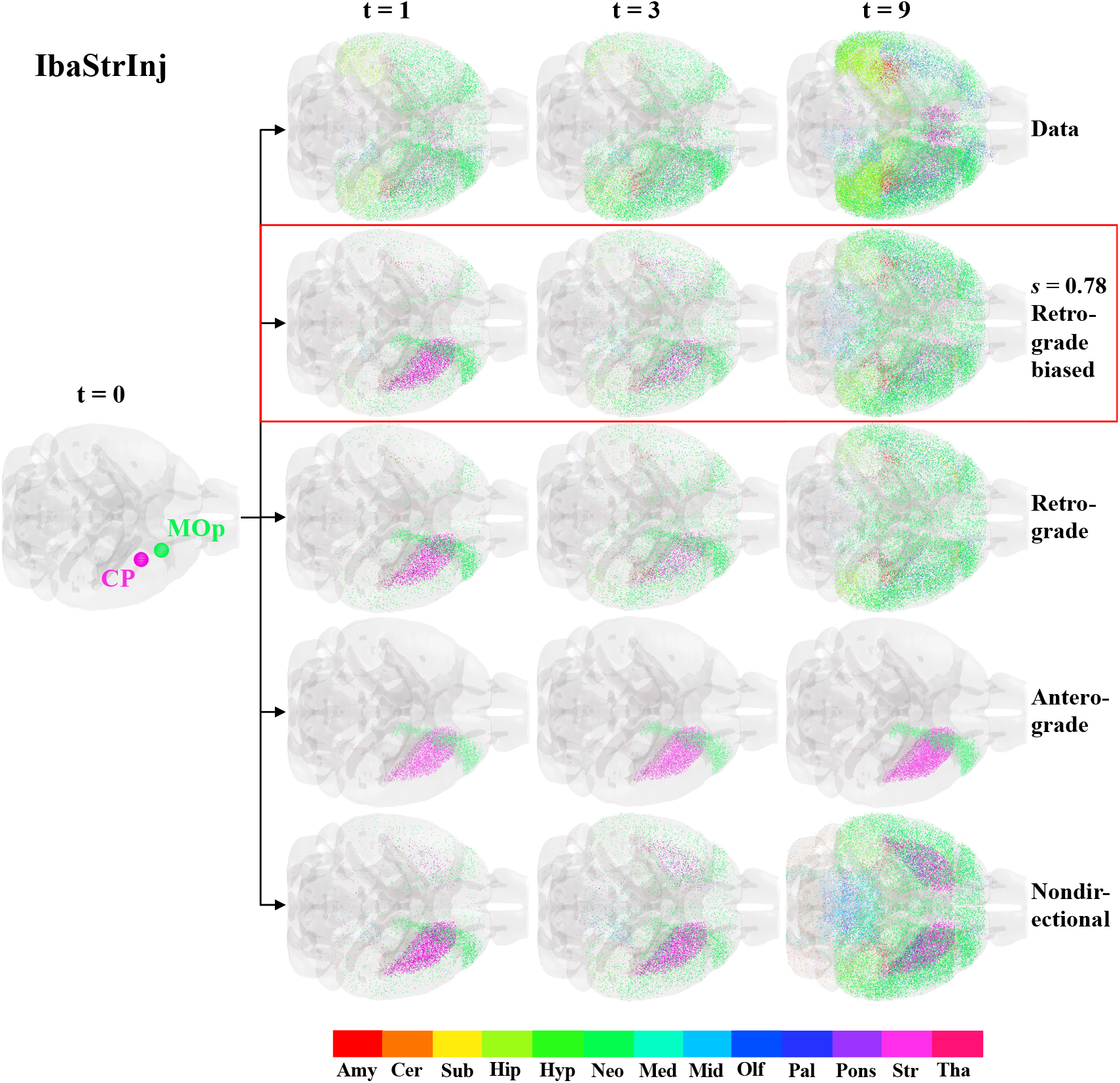
NexIS:dir models of IbaStrInj. Glass brain representations of the seed regions (CP and MOp) for the IbaStrInj experiment, visualized as spheres, and space-filling visualizations of regional tau pathology at the three quantified time points (t = 1, 3, and 9 MPI). The top row shows the observed tau distributions, with the predictions from four different NexIS:dir models shown below; in order from top to bottom: NexIS:fit-s (fitted *s* = 0.78), NexIS:ret (Ret.; *s* = 1), NexIS:ant (Ant.; *s* = 0), and NexIS:nd (N.D.; *s* = 0.5). While all models fail to capture the diffuse spreading of tau in IbaStrInj mice at 1 MPI, only by fitting the directionality bias parameter, *s*, does the model capture the distribution of tau at the last time point. There is also a clear divergence in modeled pathology by 9 MPI for different directionality biases, starting from a common seed (t = 0). Fit *s* = 0.78 indicates mixed but retrograde-biased spread. CP – caudoputamen; MOp – primary motor cortex; Amy – amygdala; Cer – cerebellum; Sub – cortical subplate; Hip – hippocampus; Hyp – hypothalamus; Neo – neocortex; Med – medulla; Mid – midbrain; Olf – olfactory; Pal – pallidum; Str – striatum; Tha – thalamus.

### NexIS:dir demonstrates that intermediate retrograde bias is the best overall model of tau spread across all studies

We extended the above analysis for IbaStrInj across all 11 mouse tauopathy models to explore if there were any trends in directionality bias. **Figure 3A** shows *R*-t curves for the four NexIS:dir models, where each colored line represents the association between end-timepoint pathology (dashed black lines; see **Figure S1** for illustrations of the observed end-timepoint tau pathology for all 11 models) and the each model’s predicted distributions across the relevant time range. We also display the optimal *s* parameters and *R* values for the NexIS:fit-s models (see also **Table S1**). While NexIS:nd and NexIS:fit-s often performed equivalently well, as exhibited by *s*_*opt*_ ≈ 0.5, NexIS:ret also exhibited good agreement and invariably outperformed NexIS:ant. As in **Figure 2**, all parameters were fit longitudinally for this analysis.

**Figure 3.**
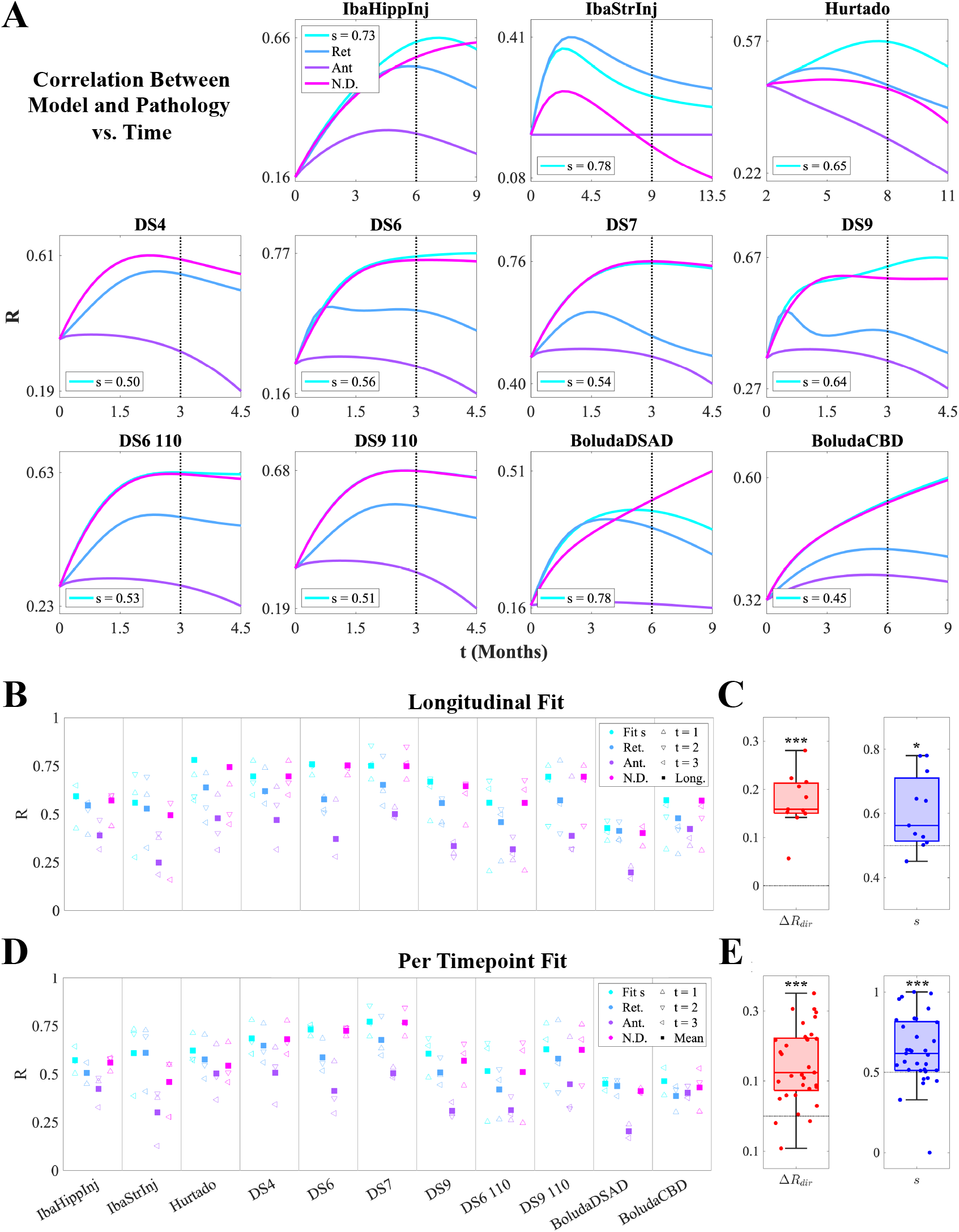
NexIS:dir model performance across all mouse experiments. **A**. Plots of Pearson’s correlation (*R*) values between the predicted tau pathology using the four types of NexIS:dir models over time, and the observed tau pathology at the last quantified time point for each of the 11 experiments (dotted vertical lines). **B**. Per-timepoint (triangles) and overall (squares) *R* values for each of the four NexIS:dir models (NexIS:fit-s, NexIS:ret, NexIS:ant, and NexIS:nd) with respect to the observed tau distributions of each of the 11 mouse experiments, where each model’s parameters were fit *longitudinally*. While NexIS:ret and NexIS:nd models often perform comparably well to NexIS:fit-s, NexIS:ant is always the worst-performing model overall. **C**. Boxplots of Δ*R*_*dir*_ (*R*_*ret*_ − *R*_*ant*_; *left*) and fitted *s* values (*right*) across experiments, assessed for longitudinally fit models. Both metrics show an overall net retrograde spread bias. **D**. Per-timepoint (triangles) and mean across time (squares) *R* values for each of the four NexIS:dir models (NexIS:fit-s, NexIS:ret, NexIS:ant, and NexIS:nd) with respect to the observed tau distributions of each of the 11 mouse experiments, where each model’s parameters were fit *to each timepoint individually*. Similar to **B**, NexIS:ant generally performs worse than all other models. **E**. Boxplots of Δ*R*_*dir*_ (*R*_*ret*_ − *R*_*ant*_; *left*) and fitted *s* values (*right*) across experiments, assessed for per-timepoint fit models. As in **C**, both metrics show an overall net retrograde spread bias. * – *p* < 0.05; ** – *p* < 0.01; *** – *p* < 0.001.

To delve into the performance in more detail, we obtained overall *R* values as well as *R* values at each quantified time point for all four models, where parameters were longitudinally fit as above (**Figure 3B**). Overall, while NexIS:fit-s is always the superior model given that it optimizes the *s* parameter, NexIS:nd is most frequently the second-best model across studies. NexIS:ret, however, outperforms it for IbaStrInj and BoludaDSAD, and as well as at certain timepoints for other studies (e.g., Hurtado). NexIS:ant is unequivocally the least predictive model across all studies. Bootstrapping analysis where we fit NexIS:fit-s to random 80%-subsets of regions showed that these results lack bias (**Figure S2**). We also assessed overall directional bias using two metrics: 1) overall Δ*R*_*dir*_ (i.e., the difference between *R*_*ret*_ and *R*_*ant*_ per study, which are plotted as blue and purple squares in **Figure 3B**, respectively); and 2) *s*_*opt*_, the optimal *s* value for NexIS:fit-s. Both metrics were statistically significantly retrograde by one-sample t-test, with Δ*R*_*dir*_ being highly statistically significant (**Figure 3C**).

We also examined an alternative fitting scheme, where we fit the *β* (spread rate) and *s* parameters at each timepoint individually after fixing *α* to its longitudinally fit value. This choice was motivated by the fact that *α*, as the global accumulation parameter, captures the increase in overall pathology burden and is best determined using all timepoints together (see **Materials and Methods**). Model performance generally followed those obtained using longitudinal fitting, with some subtle differences (**Figure 3D**). For instance, NexIS:ret exhibited a higher mean *R* across timepoints for the Hurtado study than NexIS:nd, and NexIS:ant exceeded NexIS:ret performance for BoludaCBD. However, as with the longitudinally assessed bias metrics shown in **Figure 3C**, per-timepoint Δ*R*_*dir*_ and *s*_*opt*_ also showed a highly statistically significant retrograde tendency (**Figure 3E**). Therefore, NexIS:dir conclusively demonstrated that mouse tauopathy models appear to have a net retrograde as opposed to anterograde character, with variation between individual studies.

### Spread rate, accumulation rate, and directional bias are mutually interdependent

Lastly, we postulated that the optimal values of three key NexIS:dir parameters (*α, β*, and *s*) may be correlated to each other. **Figure 4A–C** shows the linear regression between the per-timepoint fit values of each pair of parameters. Notably, *s* was negatively correlated with both *β (R*_2_ = 0.10, *p* < 0.05) and *α (R*_2_ = 0.17, *p* < 0.01) (**Figure 4A** and **B**, respectively). Of the three pairs of parameters, *α* and *β* actually showed the strongest overall correlation (**Figure 4C**, with an *R*_2_ value of 0.24 (*p* < 0.01).. We also found that there was no temporal shift in the *s* parameter when fit per-timepoint (**Figure S3**). Therefore, we conclude that only strain identity, and to some extent seeding location, contributes to net directionality.

**Figure 4.**
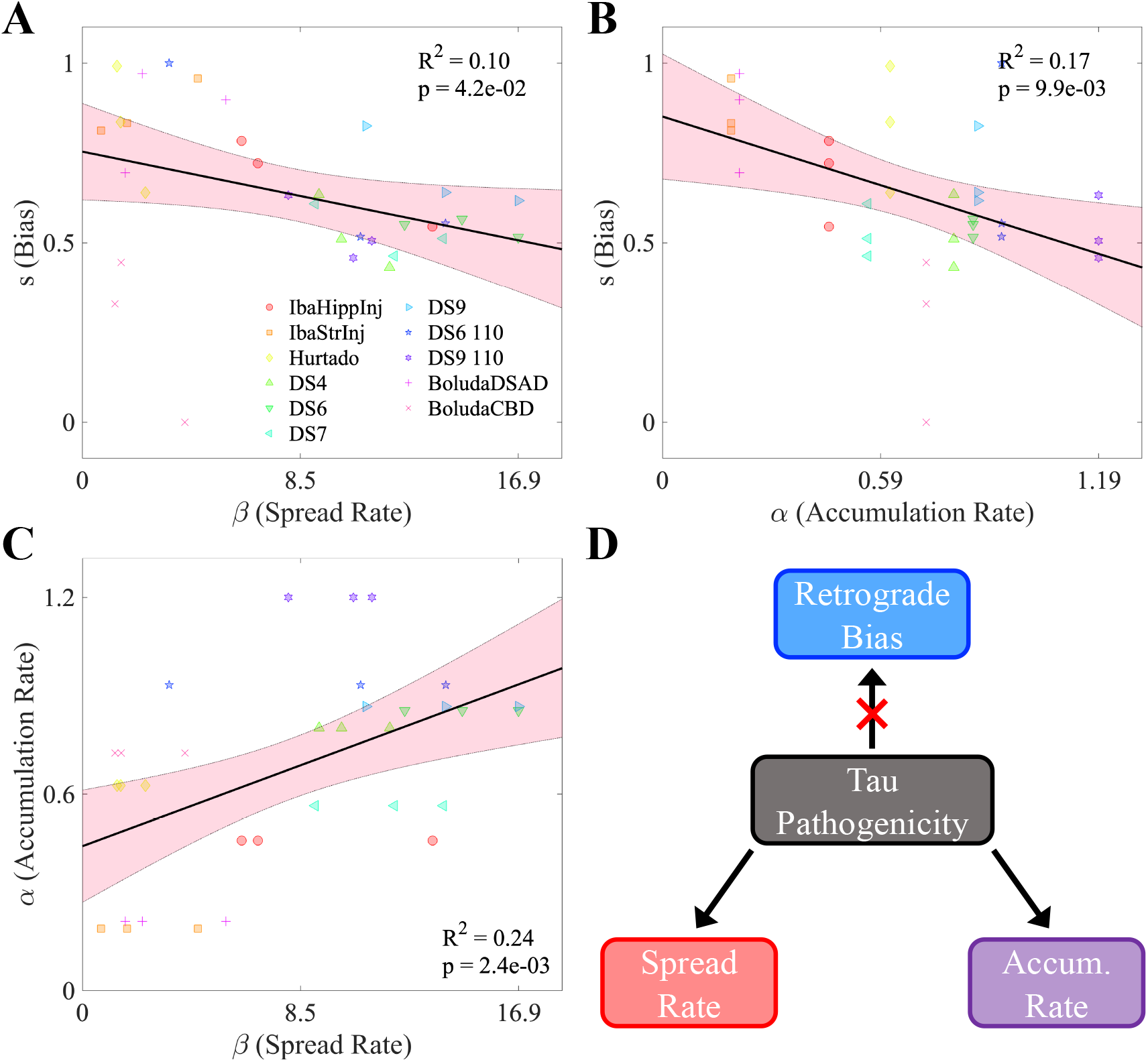
Interrelationships between NexIS:dir parameters. **A**. Scatterplot of fitted *s* values vs. spread rate parameter (*β*) values in the NexIS:fit-s model across all time points and experiments. Both parameters were fit individually per timepoint. There is a modest but statistically significant negative association between *s* and *β (p* < 0.05). **B**. Scatterplot of fitted *s* values vs. accumulation rate parameter (*α*) values in the NexIS:fit-s model across all time points and experiments. As in **A**, the *s* parameter was fit individually per timepoint, while *alpha* was fit longitudinally and fixed across timepoints (see **Materials and Methods**). There is a moderately statistically significant association between *s* and *α (p* < 0.01). **C**. Scatterplot of *α* values vs. *β* values in the NexIS:fit-s model across all time points and experiments. The parameters were fit as in **A** and **B**. There is a moderately statistically significant association between *α* and *β (p* < 0.01). **D**. Proposed model for the inter-relationships between these parameters. Overall tau pathogenicity is associated with higher accumulation and spread parameters, which are positively correlated, and is also associated with a transition from retrograde-biased to nondirectional spreading of tau along the connectome.

The implications of these findings are explored in the **Discussion** below.

## Discussion

While prior studies have firmly established the predominance of trans-synaptic spread mechanisms in driving spatiotemporal pathology development in AD ***Ahmed et al. (2014)***; ***Boluda et al. (2015)***; ***Clavaguera et al. (2009)***; ***Holmes et al. (2014)***; ***Iba et al. (2013***, 2015); ***Kaufman et al. (2017)***; ***Mezias et al. (2017)***; ***Raj et al. (2012)***, the present research is the first to explore whether the direction of trans-synaptic tau spread might be superior to non-directional spread, and whether directional bias depends on tau strain, injection site, injectate type and genetic background. We used both statistical and mathematical modeling tools to interrogate the emergence of net directional bias in tau transmission across 11 transgenic mouse datasets.

### Summary of key results

Our analyses revealed three previously unknown and striking findings. First, there is a distinct role of the polarity of axonal projections from the seeding site in where and how rapidly tau propagates into the brain, with strong statistical evidence that favors its retrograde rather than anterograde transmission. When we applied our directional network model to assess this effect on the whole brain, we found a notable net retrograde bias in network-based transmission of tau across all tauopathy studies, with anterograde-biased spread models exhibiting poor performance. Second, we observed a wide variance between individual mouse models in terms of their respective spread biases, with some exhibiting net retrograde spread and others no spread bias, thus suggesting that retrograde bias is conformation- and seeding-site-dependent. Third, extrapolating the NexIS:dir mathematical model over the whole brain network led to divergent spatiotemporal tau progression patterns that exhibited excellent agreement across all mouse models’ observed tau patterns. Therefore, the diverse spatiotemporal presentation of tauopathic conditions ***Clavaguera et al. (2013)***; ***Warren et al. (2013)***; ***Janelidze et al. (2020)***; ***Thijssen et al. (2020)***, might arise partly due to differential directional biases during trans-synaptic spread. Fourth, we found an unexpected interdependence of the NexIS:dir parameters across studies, with tau spread rate and accumulation rate being positively associated with each other and negatively associated with directional bias. This suggests that network-level phenomena may be mechanistically linked to the pathogenicity of a given tau strain. These results lead us to propose the following strain-specific model of mouse tauopathy (**Figure 4D**). First, the rates of spread and accumulation of a particular tau strain, both measures of tau “aggressiveness”, tend to be associated. By contrast, spread bias (in the retrograde direction) appears to be associated with less pathogenic tau strains.

Below we discuss these findings in the context of the literature.

### Potential mechanisms for net retrograde bias in tau propagation

The results of our statistical (**Figure 1**) and mathematical-modeling-based (**Figure 2** and **3**) analyses represent the first quantitative support for the possibility that a directional difference in the spread of tau pathology may be operative *in vivo*, or that this may have widespread consequences on the eventual pattern of progression within the brain. Most *in vivo* and *in vitro* studies indicate at least a bidirectional transmission of tau ***Dujardin et al. (2014)***; ***Iba et al. (2013)***; ***Tai et al. (2014)***; ***Wu et al. (2013)***. ***Iba et al. (2013)*** demonstrated the classic spread of preformed tau fibrils in P301S mouse model, and found that hippocampal seeding yields spread into entorhinal cortex (EC) and nearby cortices, raising the possibility that hippocampal efferents were involved. While these descriptive studies point to the plausibility of directional bias, here we present quantitative evidence across numerous experimental conditions that there is a notable, strain-specific retrograde bias in tau spreading. If confirmed by future bench studies, this finding has the potential to alter how we think about tau pathology progression.

We advance several potential mechanisms that may lead to this bias, building upon the hypothetical “molecular nexopathy” framework of degenerative diseases ***Warren et al. (2013)***. First, retrograde bias in tau pathology spread may be caused by progressive breakdown of the axon-soma barrier due to repeated exposure to hyperphosphorylated tau, leading to retrograde missorting of tau from axon to soma and dendrites ***Li et al. (2011)***. Because neuronal signaling and axon structural integrity become progressively degraded over the course of degenerative diseases ***Li et al. (2011)***, anterograde transynaptic transport likely becomes less frequent and more difficult. This would amount to a selection pressure towards misfolded tau strains that are more readily able to cross the synapse from dendritic bouton to axon terminal, thereby potentially leading to a retrograde bias across conditions. Aberrant tau conformations are also known to dysregulate energy-dependent transport of kinesin-1 ***Cuchillo-Ibanez et al. (2008)***; ***Rodríguez-Martín et al. (2013)***; ***Sherman et al. (2016)***; ***Stern et al. (2017)***, which leads to tau missorting over time and may underpin directional bias. This intriguing possibility has been explored mathematically in a two-neuron system ***Torok et al. (2021)***, but further exploration of the implications of transport breakdown and network directional bias is required to draw firm conclusions.

Retrograde transmission of tau might also be mediated by amyloid-*β* (A*β*), which is known to induce early missorting of tau from axons to dendrites ***Ballatore et al. (2007)***; ***Fein et al. (2008)***; ***Ittner et al. (2010)***; ***Takahashi et al. (2010)***. Intracerebral A*β* injections in P301L transgenic mice exacerbated hyperphosphorylation of tau and NFT formation, not at somatosensory cortical and hippocampcal injection sites, but rather in the retrogradely-connected basolateral amygdala ***Götz et al. (2001)***. This suggests that A*β*-induced damage to terminals or axons projecting to the injection site caused NFT formation in presynaptic cell bodies. While we lacked sufficient data to explore whether A*β* influences directionality in mouse tauopathy models, the fact that primary tauopathies (such as frontolobartemporal dementia, FTLD) and secondary tauopathies (such as AD) exhibit different etiologies suggests that A*β* may influence how tau spreads on the network. These explanations need not be mutually exclusive: A*β*-induced retrograde tau transmission should appear earlier in the disease course than retrograde spread caused by the breakdown of the axon-soma barrier, as amyloidopathy has been repeatedly shown to precede or co-occur with tauopathy in AD ***Bolmont et al. (2007)***; ***Götz et al. (2004)***; ***Ittner et al. (2010)***.

### Directional bias is strain-specific

It is well known that tau tangle structures within disease condition are often consistent, while they can vary considerably between disease condition ***Clavaguera et al. (2013)***. For example, when tau pathology from a human AD patient is injected into a mouse, the structure of tangle aggregates in the mice mirrors that of the patient ***Clavaguera et al. (2013)***; furthermore, comorbid A*β*/tau mouse models can reproduce Braak staging of tauopathy in the absence of a seed ***Hurtado et al. (2010)***. We found that net directional bias appears to be dependent on tau strain in a study-specific manner (**Figure 3**). While the optimal directionality bias parameter values in the NexIS:fit-s models (*s*_*opt*_) were on average retrograde across studies and timepoints, we note that there is significant variation between them (**Figure 3C** and **E**; **Table S1**).

We chose these studies in part because they all shared a P301S tau mutation, 10 of the 11 mouse models had a PS19 genetic background (the last, Hurtado, was a PS19/PDAPP bigenic mouse model), disentangling the effects of different tau strains from injection sites is more challenging. We posit that tau strain is the predominant factor here rather than seeding site based on several pieces of evidence. First, IbaHippInj and IbaStrInj, which were both obtained from the same study ***Iba et al. (2013)***, differ only in seeding locations and exhibit similar retrograde biases by *s*_*opt*_ (**Table S1**). Additionally, the 6 studies from Kaufman, *et al*. (DS4, DS6, DS6 110, DS7, DS9, and DS9 110) were all seeded in PS19 mice in the left CA1 region of the hippocampus, and therefore only differ in their injectates ***Kaufman et al. (2016)*** (**Table S1**). These all exhibit marked differences in how tau pathology evolves over time (**Figure S1**), and while most are predominantly nondirectional, DS9 exhibits retrograde character (*s*_*opt*_ = 0.64). While these data are merely suggestive, strain specificity has been noted elsewhere as a general feature underlying the diversity of tauopathic diseases ***Frost and Diamond (2010)***, as well as differences in gross tangle structure between tauopathies, as diverse as Pick’s Disease, progressive supranuclear palsy, corticobasal degeneration, and others ***Clavaguera et al. (2013)***.

What differences in these properties may explain such a difference in their preferred direction of trans-synaptic spread? Pathological tau species can bypass or move through the axon-soma barrier, which in the normal state limits retrograde misplacement of tau into somatodendritic compartment, on an isoform or strain specific basis ***Zempel et al. (2017)***. Thus, it could be that certain misfolded tau strains are more readily able to break through the barrier than others. We also found that spread and accumulation rates are inversely related to directional spread bias; therefore, more aggressive tau strains are more likely to travel nondirectionally (**Figure 4**). These observations could be explained by the known dysregulation of axonal transport by tau ***Cuchillo-Ibanez et al. (2008)***; ***Rodríguez-Martín et al. (2013)***; ***Sherman et al. (2016)***; ***Stern et al. (2017)***; however, the exact mechanistic interactions involved in these processes require further modeling and experimental studies.

### Implications

The finding that the directional bias is dependent on conformation of tau has two potential implications. First, AD heterogeneity, both in terms of disease aggressiveness ***Dujardin et al. (2020)*** and patterns of tau deposition (e.g., limbic-predominant vs. hippocampal-sparing variants) ***Murray et al. (2011)***, may be explained by the interplay between aggregation rate, spread rate, and tendency towards retrograde-biased spread. Second, it may also help explain why there is such a diversity of spatial patterning between tauopathies more broadly. These lines of reasoning are based on the observation that directional bias may lead to different areas where tau eventually accumulates, which may in turn lead to differential selective regional vulnerability of different tauopathies. Already, clinical studies confirm that the two disease hallmarks, regional tau pathology and neurodegeneration, exhibit divergent spatiotemporal development patterns between conditions, pertaining to their 3R and 4R tau isoforms ***Higuchi et al. (2002)*** and sites of tau hyperphosphorylation ***Hanger et al. (2009)***, as are the brain regions showing early disease vulnerability ***Braak and Del Tredici (2011)***; ***Burrell et al. (2016)***. Other tauopathic conditions can manifest early pathology in the orbitofrontal cortex (OFC), the amygdala (AMY), nucleus accumbens (NAcc) and caudate nuclues (CN) ***Burrell et al. (2016)***; ***Chiba et al. (2012)***. Furthermore, recent clinical evidence has identified AD-specific phosphoepitopes of tau ***Janelidze et al. (2020)***; ***Palmqvist et al. (2020)***; ***Thijssen et al. (2020)***, suggesting that amyloid comorbidity in AD may influence the ways in which tau is posttranslationally modified, which in turn leads to distinct patterns of neurodegeneration relative to primary tauopathies. The differences in spatiotemporal protein progression may eventually lead to differences in symptomatic progression. Tying these effects to underlying changes in directional bias would constitute a highly parsimonious and unifying mechanistic explanation. Future studies in both experimental and simulation systems should interrogate the sources of diversity among different neurodegenerative pathologies.

## Materials and Methods

### Datasets

The data in the present study for creating the mouse connectivity network comes from the Allen Mouse Brain Connectivity Atlas (AMBCA). This connectome was derived using viral tracing and coregistered to the Common Coordinate Framework (CCF) ***Oh et al. (2014)***; in total, there were 426 regions spanning both hemispheres in the connectome used in this study (**Figure 1C**). While there are multiple ways to quantify the amount of connectivity between regions, we chose to use *total* connectivity, rather than normalizing by volume. For all modeling analyses, the connectome was min-max normalized across all region-pairs. By convention, this 426 × 426 connectivity matrix, *C*, is oriented such that each entry *c*_*ij*_ corresponds to the connectivity *from* region *i to* region *j*.

Regional tau pathology data came from several studies that had to satisfy several criteria: 1) quantification or semi-quantification was based on immunohistochemistry (IHC); 2) the mice had to have a P301S tau mutant background to avoid conflating background endogenous tau protein type with other analyses; 3) at least 40 brain regions had to be quantified; 4) at least 3 timepoints had to be quantified. We summarize the key features of the 11 mouse models (spanning 4 studies: ***Boluda et al. (2015)***; ***Hurtado et al. (2010)***; ***Iba et al. (2013)***; ***Kaufman et al. (2016)***) in **Table S1**. Further information about each dataset or study used in the present article can be found in their respective publications.

For data derived from Boluda, *et al*., Iba, *et al*., and Kaufman, *et al*., regional tau burden was manually extracted from heat map figures displaying semi-quantified pathology regions ***Boluda et al. (2015)***; ***Iba et al. (2013)***; ***Kaufman et al. (2016)***. Data from Hurtado, *et al*. represents disease staging per region, more than quantified or semi-quantified regional pathology, but was derived from a supplementary data table ***Hurtado et al. (2010)***. Using the AIBS mouse reference at-las (https://mouse.brain-map.org/static/atlas) as an anatomic reference, we manually coregistered these regional tau values into the CCF space of the AMBCA on a per-study basis.

All datasets used for the analyses in this study are available in our NexIS GitHub repository (https://github.com/Raj-Lab-UCSF/Nexis).

### NDM and NexIS

The original Network Diffusion Model (NDM) developed by Raj, *et al*. posits that the *rate of change* in tau concentration in a given region is proportional to the tau concentration differences between all other regions with which it shares a connection ***Raj et al. (2012)***. For a two-region system, the rate of change in tau in region 1 is given by:

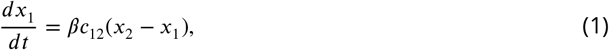

where *x*_*i*_ is the tau concentration in region *i, β* is a global spread rate parameter, and *c*_*ij*_ is the connectivity density between regions *i* and *j*. When *C* is symmetric, *c*_*ij*_ = *c*_*ji*_ and this equation represents a *graph diffusion* process. This can be generalized across all regions using the graph Laplacian, *L*, of *C*:

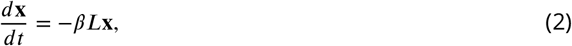

where *L* = *D* − *C* and *D* is the diagonal degree matrix of *C*. **Equation 2** is a set of ordinary, linear differential equations and has an explicit solution given by:

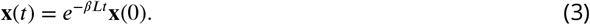

The NexIS:global model proposed by Anand, *et al*. extends this framework to account for the local accumulation of tau over time:

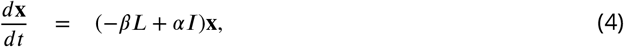

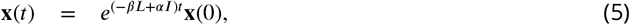

where *L* is defined as above and *I* is the identity matrix. Here accumulation is parameterized by the accumulation rate *α* ≥ 0. The above equation therefore models an exponential growth of pathology as a function of local tau concentration; the nonnegativity constraint is justified by the fact that all 11 studies exhibit an increase in overall tau burden over time. In this study we require the accumulation rate to be the same everywhere in the brain – assuming no region-intrinsic effects – hence *α* is a global rate parameter.

### NexIS:dir

The above framework was proposed and implemented for symmetric connectomes only; that is, when *c*_*ij*_ ≡ *c*_*ji*_. This is particularly appropriate for network modeling of human pathology, since human connectomes are obtained using diffusion tensor imaging (DTI), which is unable to distinguish the polarity of white matter tracts in the brain. However, the AMBCA is fully directed, and therefore it need not be assumed that flows along connection *c*_*ij*_ are equivalent to those along *c*_*ji*_. Here, we define two extreme cases for NexIS:dir: 1) *purely retrograde spread*, where *positive flow* from region *j* to region *i* occurs along *c*_*ij*_ ; 2) *purely retrograde spread*, where *positive flow* from region *j* to region *i* occurs along *c*_*ji*_. Mathematically, the two-region retrograde case is given by:

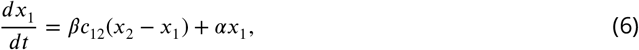

and the two-region anterograde case is given by:

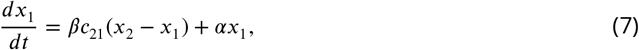

As above, these generalize across the network using the graph Laplacian (**Equation 4**); however, the form of the Laplacian differs between these two cases:

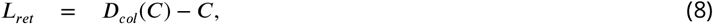

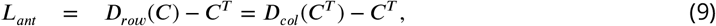

where *D*_*row*_ and *D*_*col*_ are the row and column degree operators, respectively. *D*_*row*_(*C*) produces a matrix where each diagonal entry is the *outgoing* degree of *C* and *D*_*col*_(*C*) produces a matrix where each diagonal entry is the *incoming* degree of *C*.

With NexIS:dir, we propose that directionally biased spread need not be *entirely* aligned in the anterograde or retrograde directions as in **Equations 8** and **9**. To accomplish this, we first define a new *directionally weighted* connectome, *C*_*s*_, as follows:

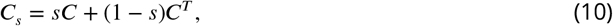

Here, *s* is a proportionality constant weighting the contributions of *C* and *C*^*T*^ to the flows on the network, and is bounded between 0 and 1. It is clear by inspection that *C*_1_ = *C* and *C*_0_ = *C*^*T*^. The directional Laplacian, then, is simply:

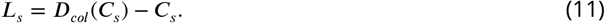

Again, *L*_1_ = *L*_*ret*_ and *L*_0_ = *L*_*ant*_. Finally, the full NexIS:dir model is given by:

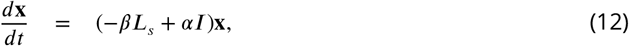

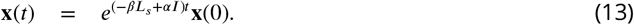

We note one other special case of NexIS:dir: the case when *s* = 0.5. *C*_0.5_ = 0.5(*C* +*C*_*T*_), and therefore *c*_0.5,*ij*_ = 0.5(*c*_*ij*_ + *cji*) = *c*_0.5,*ji*_ and *C*_0.5_ is symmetric. Therefore, we define the nondirectional Laplacian *L*_*nd*_ = *L*_0.5_, which is equivalent to the NexIS:global case (**Equation 4**).

We use NexIS:fit-s, NexIS:ret, NexIS:ant, and NexIS:nd throughout this study, where these are distinguished by the forms of the Laplacian that they use in **Equation 12** (*L*_*s*_, *L*_*ret*_, *L*_*ant*_, and *L*_*nd*_, respectively).

### Parameter fitting

We fit all parameters of the NexIS:dir model using MATLAB’s fmincon nonlinear optimization algorithm for each study individually. We utilized two different fitting schemes: 1) longitudinal fitting, where all available timepoints were fit simultaneously; and 2) per-timepoint fitting, where *α* was first fixed to its longitudinally fit value and then the remaining parameters were fit to each time-point individually. There is a latent parameter, *γ*, representing the proportionality constant relating amount of tau injected into seeded regions and the tau density as quantified by IHC; this scale factor varies by experimental condition and is necessary for making any fit of *α* meaningful. It also prevents the fitting of *α* in a per-timepoint fashion, as there is an identifiability problem between *α* and *γ* when fitting only one timepoint from seed. Therefore, for per-timepoint fitting, *γ* was also fixed alongside *α*.

The regional data space differs from the regional connectome space (i.e., the CCF) and between studies; notably, not all regions of the CCF were quantified in any given study. We therefore simulated the spread of pathology on the CCF atlas, producing 426 × number-of-timepoint matrices of NexIS:dir predictions. To optimize the parameters for a given mouse model, we first coregistered these predictions into that model’s specific data space and then assessed the cost. The cost function used was the Lin’s Concordance Correlation Coefficient, as previously described ***Lin (1989)***; ***Anand et al. (2022)***.

### Statistical analysis

All statistical analysis involved performing standard linear regressions, Pearson’s correlation, and t-tests using in-built MATLAB functions. *p*-values were subjected to the Bonferroni correction where appropriate.

### Visualization

All analysis figures were produced using MATLAB plotting functions. The glass brain visualizations were generated using an in-house tool, Brainframe (https://github.com/Raj-Lab-UCSF/Brainframe).

## Code and data availability

The data and code for these analyses and figures can be found in the NexIS repository (https://github.com/Raj-Lab-UCSF/Nexis).

## Acknowledgments

We would like to acknowledge the Allen Institute for Brain Science and in particular their mouse reference and connectivity atlas toolboxes and research teams. Without their exemplary contributions to neuroscience and neuroanatomy, the present work would not be possible. We would also like to acknowledge Pedro D. Maia of the University of Texas at Arlington, who was instrumental in the development of the NexIS model. This research was supported by the following grants from the National Institutes of Health: R01NS092802, R01EB022717, RF1AG062196, R56AG064873.

## Author Contributions

JT wrote the analysis code in MATLAB. JT and AR designed the models and co-wrote the manuscript. CM performed the literature searches to find the previously published datasets or tables and figures used as the “empirical data” here, performed the analyses to collect results and designed the figures and tables.

## Compliance with ethical standards

Conflict of interest: The authors declare no competing financial interests. Ethical approval and consent to participate: All mouse-work datasets in the present work was obtained from datasets originally presented in or with other already published work already declaring that all animal work was done under approved protocols and animal care standards.

