## Supplemental Figures and Tables for "Directionality bias underpins divergent spatiotemporal progression of Alzheimer-related tauopathy in mouse models"

| Name | Mouse Model Description | $R_{\text{fit-s}}$ | $s_{opt}$ |
|---|---|---|---|
| **IbaStrInj** [1] | PS19 mice seeded with synthetic PFFs from 2N4R P301S tau (T40/PS) and from truncated P301L tau (K18/PL) in the right CP and right MOp | $\mathbf{0.56}^{***}$ | 0.78 |
| **IbaHippInj** [1] | PS19 mice seeded with synthetic PFFs from 2N4R P301S tau (T40/PS) and from truncated P301L tau (K18/PL) in the right DG | $\mathbf{0.59}^{***}$ | 0.73 |
| **Hurtado** [2] | Unseeded doubly transgenic PS19/PDAPP mouse | $\mathbf{0.78}^{***}$ | 0.65 |
| **DS4** [3] | PS19 mice seeded with tau from AD brain homogenate in the left CA1 area | $\mathbf{0.70}^{***}$ | 0.50 |
| **DS6** [3] | PS19 mice seeded with P301S mouse-derived, fibril-like cytoplasmic inclusions ("threads") in the left CA1 area | $\mathbf{0.76}^{***}$ | 0.56 |
| **DS6 110** [3] | PS19 mice seeded with a 1:10 dilution of the **DS6** tau strain in the left CA1 area | $\mathbf{0.56}^{***}$ | 0.53 |
| **DS7** [3] | PS19 mice seeded with recombinant fibrils with prominent nuclear inclusions ("speckles") in the left CA1 area | $\mathbf{0.75}^{***}$ | 0.54 |
| **DS9** [3] | PS19 mice seeded with recombinant fibrils with prominent nuclear inclusions ("speckles") in the left CA1 area | $\mathbf{0.67}^{***}$ | 0.64 |
| **DS9 110** [3] | PS19 mice seeded with a 1:10 dilution of the **DS9** tau strain in the left CA1 area | $\mathbf{0.69}^{***}$ | 0.51 |
| **BoludaDSAD** [4] | PS19 mice injected with DSAD brain homogenate in the left CA1 and left primary somatosensory areas (LH) | $\mathbf{0.43}^{***}$ | 0.78 |
| **BoludaCBD** [4] | PS19 mice injected with CBD brain homogenate in the left CA1 and left primary somatosensory areas (LH) | $\mathbf{0.57}^{***}$ | 0.45 |

**Table S1: Summary of key NexIS:directed results**. List of the tauopathy datasets explored here with descriptions of each experiment's mouse genetic background, injection site, and type of tau injected. The Pearson's correlation values for the directionality-fitted models ($R_{\text{fit-s}}$) and the optimal directionality parameter ($s_{opt}$) values are provided. All studies were quantified tau pathology within hemispheres ipsilateral and contralateral to the injection site separately with the exception of Hurtado, which was bilaterally averaged. PFF – preformed fibrils; DSAD – Down Syndrome Alzheimer's disease; CBD – corticobasal dengeneration. * – $p < 0.05$; ** – $p < 0.01$; *** – $p < 0.001$.

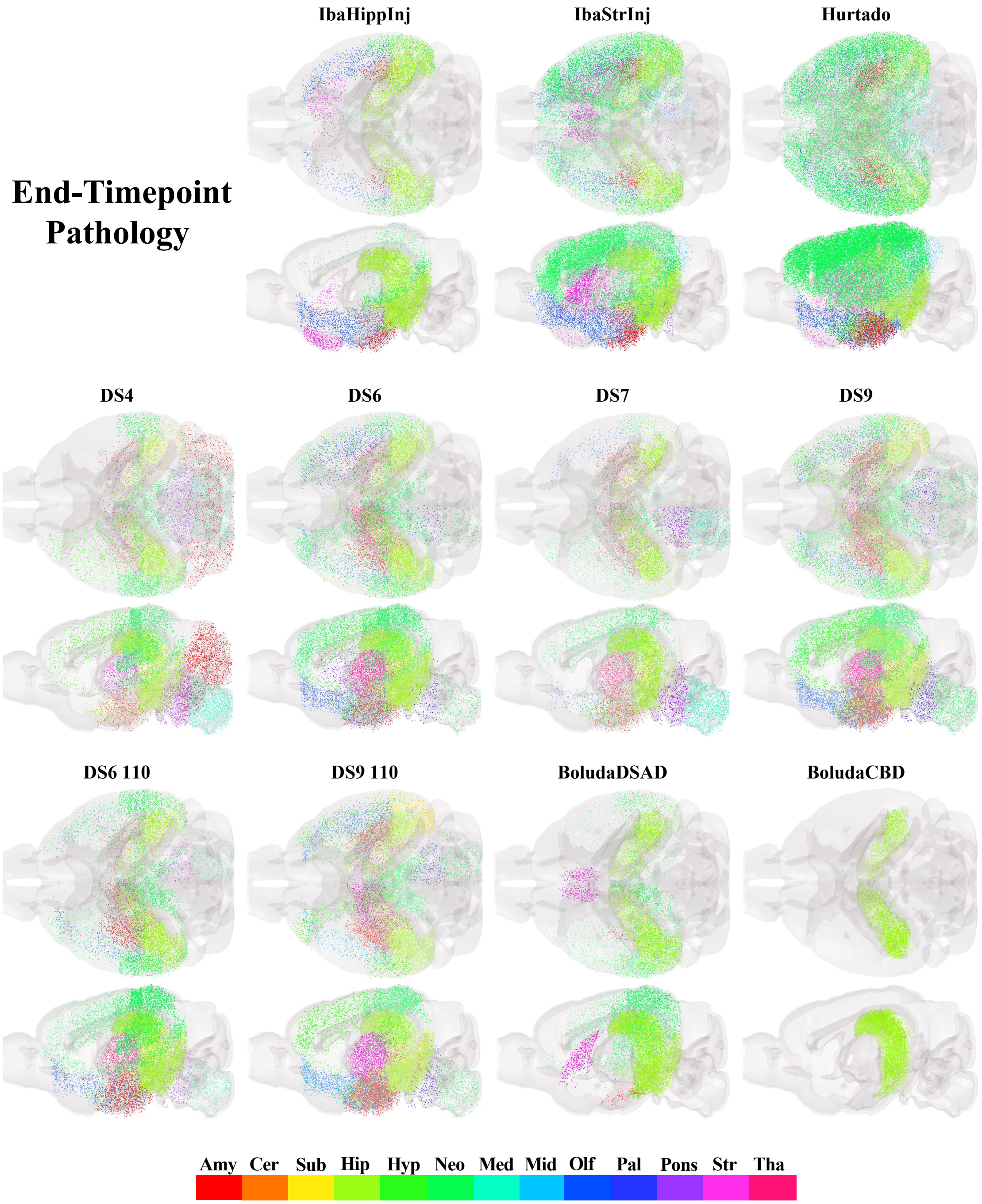


**Figure S1: End-timepoint pathology glass brains.** End-timepoint pathology for each of the twelve mouse tauopathy datasets, plotted in axial and sagittal views. See **Table 1** for descriptions of these datasets.

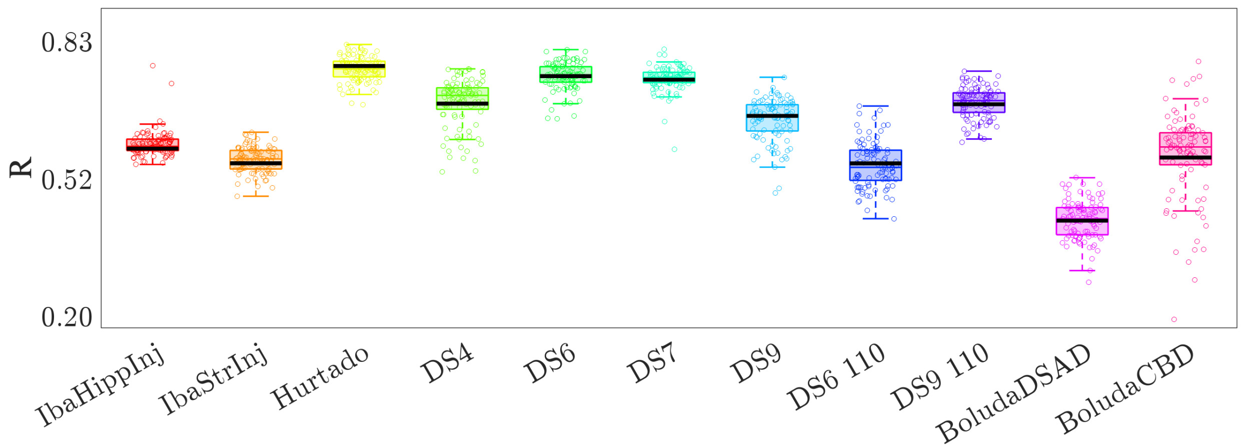


**Figure S2: Bootstrapping analysis reveals no fit bias** Boxplots and scatter representing the Pearson's R values obtained when NexIS:fit-s was fit on 100 random subsets of 80% of brain regions, alongside the fits to all regions (black lines). The fitting to all regions does not appear to introduce strong bias. See **Table 1** for descriptions of these datasets and **Methods** for more details.

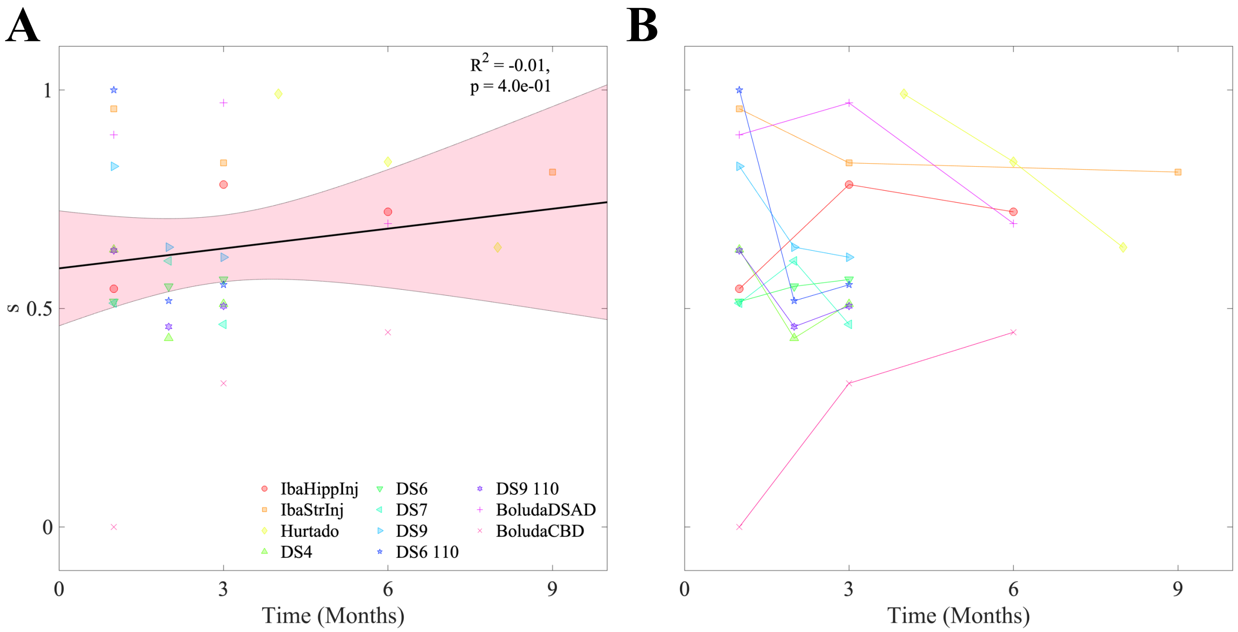


**Figure S3: There is no net temporal relationship across studies with respect to directionality bias A.** Linear regression reveals that per-timepoint-fit $s$ values do exhibit a shift over time across mouse models. **B.** The same data as **A**, plotted as trajectories for individual mouse models, showing that there is no consistent trend within studies. See **Table S1** for descriptions of these datasets.
